# 16S rRNA:rDNA ratios and cell activity staining reveal consistent patterns of soil microbial activity

**DOI:** 10.1101/435925

**Authors:** Alan W. Bowsher, Patrick J. Kearns, Ashley Shade

**Author notes:** Address correspondence to Dr. Ashley Shade,. Department of Biology, Tufts University, Medford, Massachusetts, USA.

## Abstract

Microbial activity plays a major role in the processes that support life on Earth. Nevertheless, across diverse ecosystems many microbes are in a state of dormancy, characterized by strongly reduced metabolic rates. Of the methods used to assess microbial activity-dormancy dynamics, 16S rRNA: rDNA amplicons (“16S ratios”) and active cell staining with 5-cyano-2,3-ditolyl tetrazolium chloride (CTC) are two of the most common, yet each method has its own limitations. To better understand the applicability and potential complementarity of these two methods, we conducted two experiments investigating microbial activity in the rhizosphere. In the first experiment, we treated corn rhizosphere soil with common phytohormones to simulate plant-soil signaling during plant stress, and in the second experiment, we used bean exposed to drought or nutrient enrichment to more directly assess the impacts of plant stress on soil microbial activity. Overall, 16S ratios revealed numerous taxa with detectable RNA but no detectable DNA. However, overarching patterns in percent activity across treatments were unaffected by the method used to account for active taxa, or by the threshold 16S ratio used for taxa to be classified as active. 16S ratio distributions were highly similar across microbial phyla and were only weakly correlated with ribosomal operon number. Lastly, over relatively short time courses, 16S ratios are responsive earlier than CTC staining, a finding potentially related to the temporal sensitivity of activity changes detectable by the two methods. Our results suggest that 16S ratios and CTC staining provide robust and complementary estimates of bulk community activity.

**Importance:** Although the majority of microorganisms in natural ecosystems are dormant, relatively little is known about the dynamics of the active and dormant microbial pools through both space and time. The limited knowledge of microbial activity-dormancy dynamics is in part due to uncertainty in the methods currently used to quantify active taxa. Here, we directly compared two of the most common methods (16S ratios and active cell staining) for estimating microbial activity in rhizosphere soil, and found that they were largely in agreement in the overarching patterns, suggesting that either method is robust for assessing comparative activity dynamics. Thus, our results suggest that 16S ratios and active cell staining provide robust and complementary information for measuring and interpreting microbial activity-dormancy dynamics in soils. They also support that 16S rRNA:rDNA ratios have comparative value and offer a high-throughput, sequencing-based option for understanding relative changes in microbiome activity.

## Introduction

Microbial activity plays a fundamental role in the processes that support life on Earth (1), influencing global carbon and nutrient cycling (2, 3), atmospheric composition (4), and ecosystem productivity (5). Given these essential global-scale functions, it is perhaps surprising that active microbes (those that are growing or reproducing, or those that respond relatively quickly to substrate input) represent a very small proportion of total microbial community (reviewed in (6)). Across diverse ecosystems, the majority of the total microbial community is in a state of dormancy, characterized by strongly reduced metabolic rates and a slow response to substrate input (6, 7). Dormancy is a key contributor to the maintenance of microbial diversity by allowing microorganisms to persist during unfavorable environmental conditions (8). Although dormancy initiation and resuscitation to an active state have ecological and evolutionary consequences (7–9) with implications for ecosystem function (10, 11), we know little about the dynamics of the active and dormant microbial pools through both space and time. Investigations of the causes and consequences of microbial activity-dormancy dynamics are needed to better understand microbial community structural and functional resilience, and to advance goals to predict microbial responses to global change (12).

The limited knowledge of microbial activity-dormancy dynamics is in part due to uncertainty in the methods currently used to quantify active taxa (6). One of the most common approaches is the use of 16S ribosomal rRNA sequencing. Given the relatively short half-life of ribosomal RNA, the presence of 16S ribosomal transcripts (hereafter “rRNA) is generally assumed to indicate recent metabolic activity, and numerous studies have used rRNA to characterize growing or active communities (reviewed in (13)). In particular, pairing both 16S ribosomal transcripts (rRNA) as well as the 16S ribosomal gene (rDNA) sequencing allows for calculation of 16S rRNA:rDNA ratios, which attempts to normalize rRNA levels by the abundance of that taxon in the community and quantify its relative level of activity (8, 11, 14, 15). Taxa with 16S ratios greater than a specified threshold are considered ‘active’, and most studies report using a threshold of 1.0, which indicates more rRNA reads than rDNA reads for those taxa (6, 8, 16). One limitation to this approach is that using an arbitrary threshold to distinguish active from dormant taxa may be problematic in diverse communities (13, 17), given that rRNA production and growth rate are not necessarily always correlated (24–29). Another challenge in assessing 16S ratios is the occurrence of ‘phantom taxa’: taxa that are detected in rRNA but not rDNA sequences (18) which leads to a zero-denominator (and thus undefined) 16S ratio. Phantom taxa are unexpected in sequencing datasets, since the presence of rRNA necessitates the presence of the template rDNA from which it was transcribed; yet nearly 30% of the OTUs detected in a recent study of atmospheric samples were phantoms (18). The authors of that study concluded that the prevalence of phantom taxa were driven largely by rare taxa (i.e. those with low abundance in rDNA sequences) exhibiting disproportionately high activity compared to more abundant taxa; thus phantom taxa arise largely due to sampling stochasticity (18), or alternatively, by in-process transitions from rarity to prevalence (i.e., conditionally rare taxa, (19)). Although these considerations have led researchers to suggest that 16S ratios may be best interpreted as ‘potential microbial activity’, 16S ratios have nevertheless been used to inform microbial activity-dormancy dynamics in a broad range of ecosystems (11, 14, 15, 18).

In addition to 16S RNA/DNA sequencing, a variety of other methods are currently used for distinguishing active microbes. These include staining with tetrazolium salts, which are reduced by dehydrogenases in active cells to fluorescent products which can be visualized and measured ((20) and references therein), stable isotope probing to quantify uptake of substrates or water (21), and meta-transcriptomics to determine changes in functional gene transcripts following experimental perturbation (22). Of these methods, active cell staining, primarily with the activity stain 5-cyano-2,3-ditolyl tetrazolium chloride (CTC), is a common approach because it is economical and executable without specialized equipment. Actively respiring cells convert CTC to an insoluble red fluorescent formazan product, which can be visualized by fluorescence microscopy (23). In addition, CTC staining can be coupled with a general DNA stain to compare active and total cell counts in a microbial community, allowing for determination of percent activity (15, 24). As with the 16S ratio method for determining microbial activity, the CTC staining method has several limitations which can affect its interpretation. For example, CTC staining can be toxic to some bacterial species (20, 25), and not all actively respiring strains are able to take up the stain efficiently (20, 24), potentially leading CTC staining to underestimate the true proportion of active cells in the community (26). Despite these potential caveats, CTC staining remains a popular method for analyses of microbial activity in a broad range of environmental samples (7, 15, 24).

Although the 16S ratio method and the CTC staining method are both commonly used in investigations of microbial activity, little comparative work has been done to assess the level of agreement between the two methods. One of the few studies to use both methods to assess microbial activity found that the active portion of the community was between 1.5-and 5-fold higher when using 16S ratios versus CTC staining in microcosms of estuarine water samples (15). One potential reason for the disagreement between the 16S ratio method and the CTC staining method is not only that the two methods present different biases as described above, but that they measure two different things: the 16S ratio method is used to assess whether a particular taxon is active, while the CTC staining method is used to assess whether a given cell is active. Importantly, there are situations in which we might expect the proportion of active taxa and the proportion of active cells in a community to be very different, such as in communities in which rare taxa are disproportionately active compared to abundant taxa (15, 18, 27). Therefore, although the two methods may not always produce similar estimates of microbial activity-dormancy dynamics, both inform on fundamental aspects of microbial communities. Additional studies directly comparing 16S ratio and CTC staining is needed in order to better understand the level of agreement, and the potential complementarity, of the two methods in assessing microbial activity.

Here, our objectives were to explore factors underlying the calculation and interpretation of 16S ratios, and to directly compare estimates of activity of microbial communities using 16S ratios and CTC-based cell staining. We conducted two separate experiments analyzing microbial activity in plant rhizosphere soil. Given that the rhizosphere is a highly dynamic microbial system in which plants can influence both microbial community structure and function (28, 29), we considered rhizosphere soil to be particularly relevant for analyses of microbial activity and foundationally important for understanding plant-microbe-soil feedbacks. First, we conducted a laboratory microcosm experiment using soil collected from the rhizosphere of corn, and treated the soil with several plant phytohormones to assess the impacts of common plant stress signals on soil microbial activity. Second, we grew bean plants under either drought or nutrient-enriched conditions to more directly assess the impacts of plant stress on soil microbial activity. Specifically, we asked the following questions: 1) for 16S ratio-based studies, what is the extent of phantom taxa, and how does the handling of these taxa influence estimates of microbial activity and patterns across treatments? 2) How does the threshold for quantifying ‘active’ taxa influence patterns across treatments? 3) How do divergent traits such as 16S rRNA operon copy number impact the distribution of 16S ratios within and across phyla? 4) Do 16S ratio and CTC-based staining methods produce similar estimates and/or patterns of microbial activity across diverse soil treatments?

## Results and Discussion

We conducted two separate experiments in rhizosphere soils under a variety of treatment conditions in order to explore the generality of our findings. In the first experiment, we collected soil from a long-term agricultural research field in which corn (*Zea mays* L.) had been planted for eight consecutive years. In the laboratory, we exposed the soil to several pre-treatments: ‘pre-dry’ (soil was sampled before any treatments were initiated), ‘post-dry’ (soil was dried for three days and then sampled), and ‘post-water’ (soil was partially re-wetted, allowed to acclimate for six days, and then sampled). Next, soil replicates were treated with either abscisic acid (ABA), indole-3-acetic acid (IAA), jasmonic acid (JA), or salicylic acid (SA), or water as a control, and sampled after 24 hours. Thus, the corn experiment was designed to assess the impacts of several different abiotic/biotic stresses on rhizosphere soil, including soil drying and re-wetting, as well as the application of common plant stress phytohormones, which can be exuded by plant roots under a variety of stress conditions (30). In the second experiment, we grew common bean (*Phaseolus vulgaris* L. cv. Red Hawk) in agricultural field soil in a controlled-environment growth chamber. Replicate plants were exposed to either drought (water-withholding) or nutrient enrichment (additional fertilizer) treatments compared to control plants, then rhizosphere soils were sampled after five weeks of plant growth. Thus, this experiment was designed to more directly assess the impacts of plant stress on analyses of soil microbial activity in the rhizosphere. Overall, we anticipated that the differential treatment conditions within and across the corn and bean soil experiments, as well as the presence of actively growing plants continuously providing labile carbon to the soil microbial communities in the bean but not the corn soil study, would inform on the broad applicability of the 16S and CTC staining methods for assessing microbial activity in diverse study systems.

### ‘Phantom taxa’ are common and persistent in diverse soil treatments

A prerequisite for assessing microbial activity from 16S ratios is determining how to handle ‘phantom taxa’: OTUs that are detected in the RNA community but not in the DNA community of a given sample (18). Such taxa produce undefined 16S RNA:DNA ratios due to a denominator of zero, eliminating those taxa from the dataset. Therefore, we assessed the prevalence of phantom taxa (taxa with RNA reads > 0 and DNA reads = 0 in a given sample) in both the corn and bean rhizosphere soil datasets. We also assessed the prevalence of ‘singleton’ phantom taxa (taxa with RNA reads = 1 and DNA reads = 0 in a given sample), given that such taxa are particularly ambiguous in terms of activity. We repeated these analyses across a range of sequencing depths per sample, given that a recent study reported that sampling stochasticity was a key factor driving the occurrence of phantom taxa (18). Across a range of subsampling levels (using a step-size of 5000 reads per sample), we found that phantom taxa comprised between 5% and 60% of the total OTUs across both the corn and the bean rhizosphere soil datasets (Fig. 1A and B, Fig. S1A and B). Similarly, ‘singleton’ phantom taxa were fairly common (1-35% and 2-25% of the total OTUs in the corn and bean soil datasets, respectively) (Fig. 1C and D, Fig. S1C and D). The reader should note that the sample size for each treatment generally decreased as sequencing depth increased because samples are excluded when their total read count is less than a given sequencing depth (Fig. S2). The relatively high prevalence of phantom taxa is in agreement with a recent study of atmospheric samples (18), and suggests that rare taxa are relatively more active than abundant taxa, resulting in their presence in RNA but not DNA sequences. This conclusion is supported by the relatively small influence of increasing sequencing depth on the occurrence of phantom taxa (i.e. increased sequencing only minimally increases the detection of extremely rare taxa).

**Fig 1.**
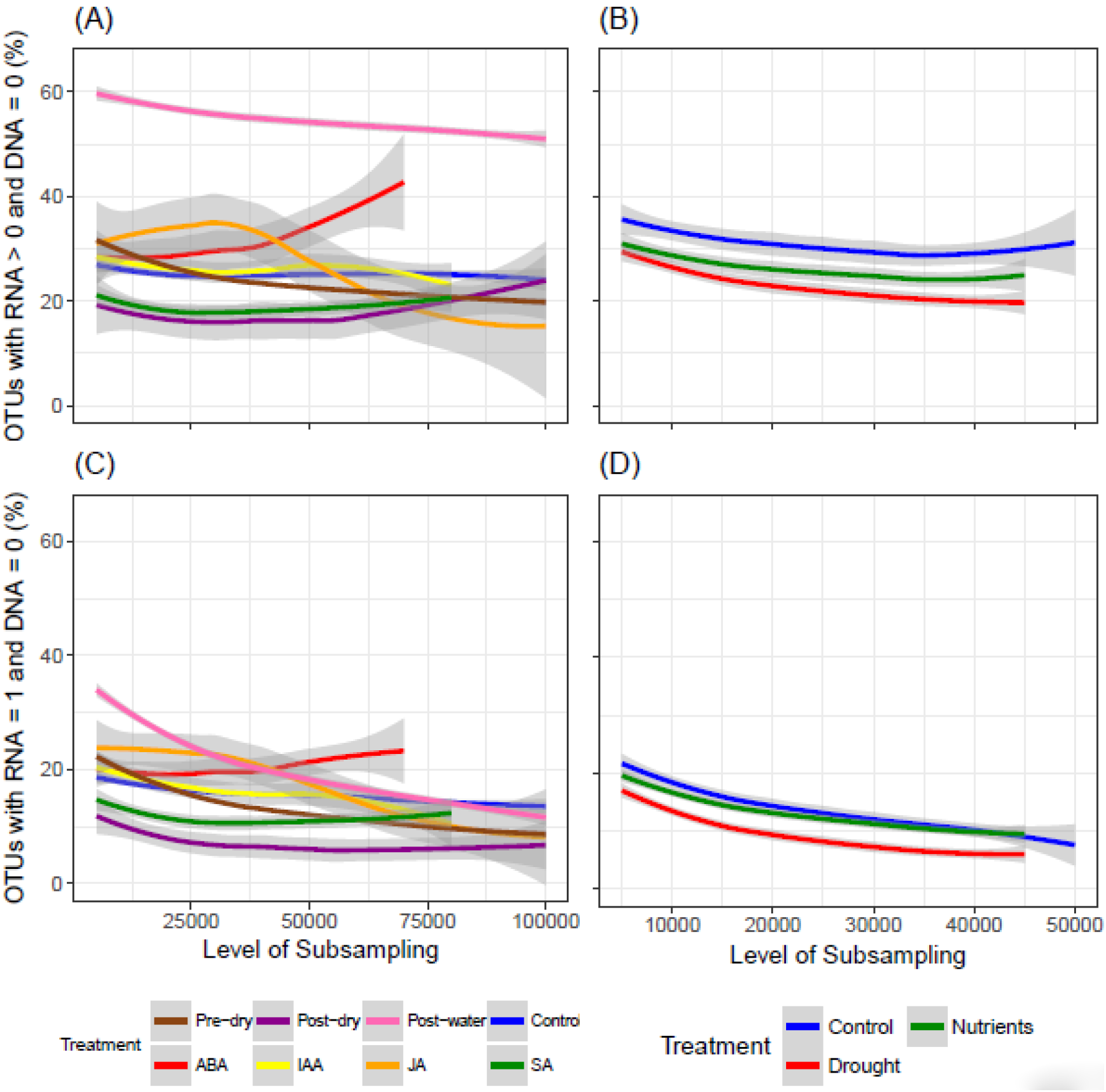
Prevalence of taxa with 16S RNA reads but zero 16S DNA reads (A, B) (i.e. ‘phantom taxa), or a single 16S RNA read and zero 16S DNA reads (C, D) in rhizosphere soil of corn (A,C) and bean (B,D) as a function of sequencing subsampling level. Shown are best fit lines using the loess smoothing function (see Supplementary Figure S1 for same plots but including individual data points). Gray shading around the smoothing lines are 95% confidence intervals.

### Distinct methods for handling phantom taxa lead to similar ecological patterns of microbial activity

Given the prevalence and persistence (i.e., their occurrence regardless of sampling depth) of phantom taxa in our sequencing datasets, our next aim was to establish how to handle phantom taxa for calculation of 16S ratios. We compared four different methods for calculating 16S ratios in the presence of phantom taxa, which we refer to here as Methods 1, 2, 3, and 4 for simplicity (Fig. 2). In both the corn and the bean rhizosphere soil datasets, all four methods for calculating 16S ratios produced similar patterns across treatments (Fig. 3A and B). In corn soil, percent activity sharply decreased from the pre-dry to the post-dry treatment, then sharply increased from the post-dry to the post-water treatment (Fig. 3A). In addition, although increasing the threshold 16S ratio for defining taxa as ‘active’ generally reduced the magnitude of treatment effects on microbial activity, the large impact of the post-water treatment on microbial activity was apparent even at a threshold ratio of five. Similar analyses of ratio thresholds have been performed in in other studies (7, 8), which collectively suggest that even conservative ratio thresholds provide robust ecological patterns. On the other hand, although percent activity in the bean experiment was generally higher in the control than in the drought or nutrient treatments across Methods 1-4, the magnitude of this difference decreased as threshold 16S ratio increased (Fig. 3B). Differences among treatments disappeared when compared at a threshold 16S ratio of five, indicating a relatively narrow window for capturing differences in microbial activity in the bean experiment. Nevertheless, a recent review suggests that most studies use a threshold 16S ratio between 0.5-2 to determine active taxa (6), indicating that a threshold of five may simply provide an overly-conservative view of activity in microbial communities.

**Fig 2.**
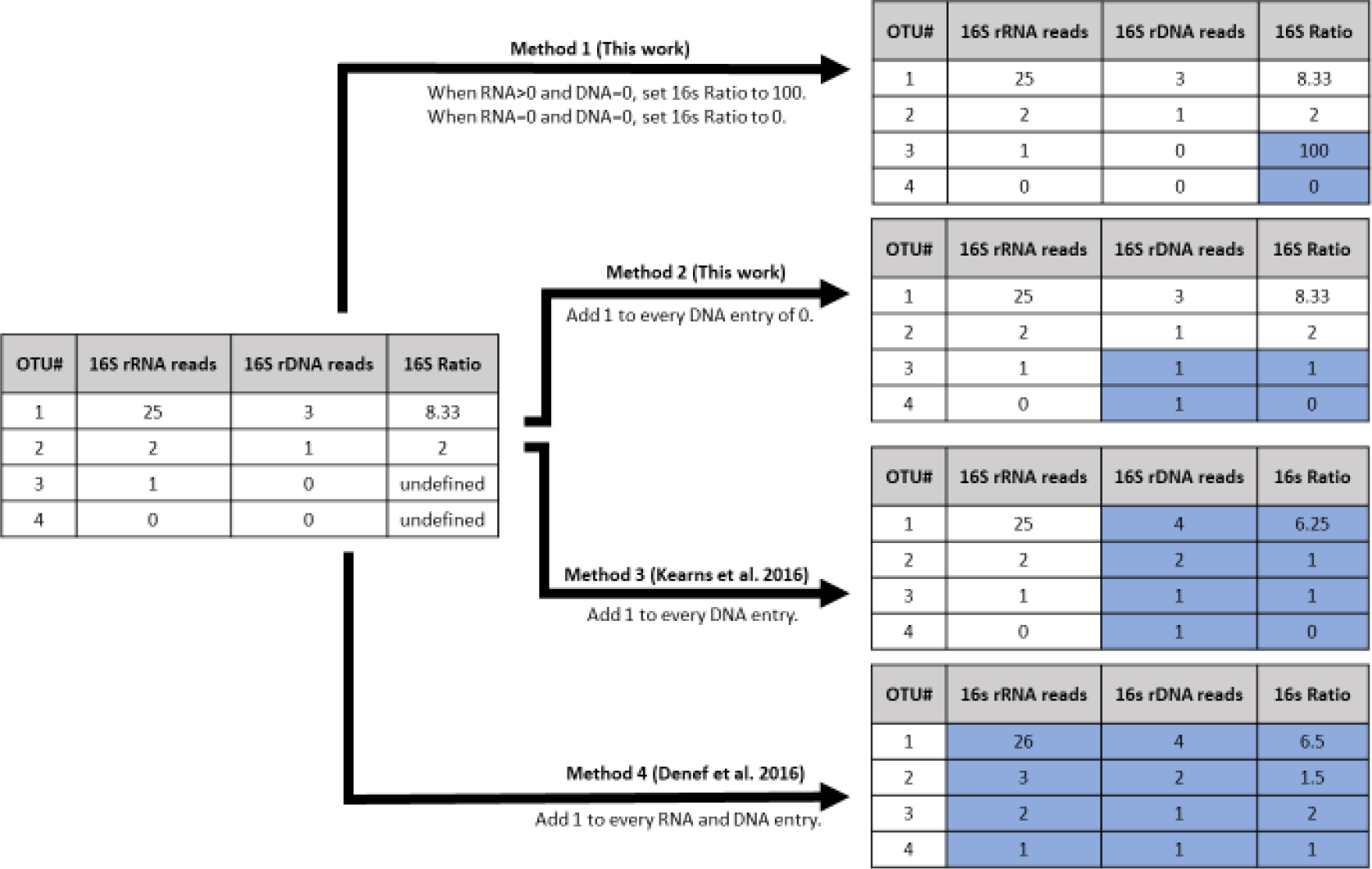
Conceptual diagram depicting the impacts of four distinct methods for calculating 16S rRNA:rDNA ratios in the presence of ‘phantom taxa’ (i.e., OTUs in a given sample with 16S RNA reads but zero 16S DNA reads, which always produce undefined 16S ratios due to a zero denominator). The input OTU table for a given sample along with 16S ratios is shown on the left, while the resulting OTU tables and 16S ratios are depicted on the right (changes are shaded blue). Four different sequencing scenarios in a hypothetical sample are considered: OTU1, in which the number of RNA reads is much larger than the number of DNA reads but both are present; OTU2, in which the number of RNA and DNA reads are both low but present; OTU3, in which the number of RNA reads is one and the number of DNA reads is zero; and OTU4, in which the number of both RNA and DNA reads is zero.

**Fig 3.**
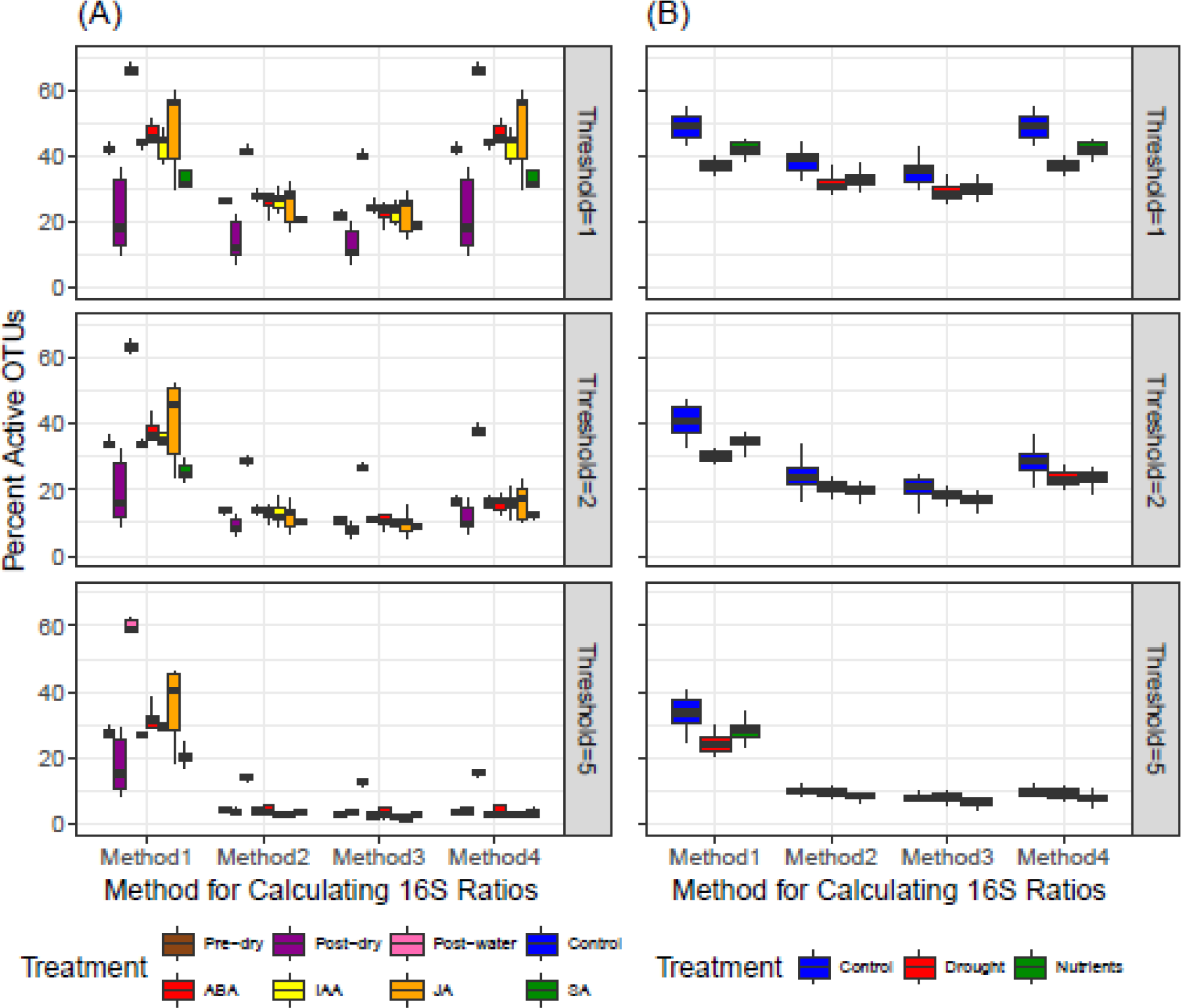
Comparison of the proportion of taxa that are active (i.e. percentage of total OTUs with 16S rRNA:rDNA ratio greater than a given threshold) in rhizosphere soil of bean (A) and corn (B) following each of four methods for calculating 16S ratios. See Figure 2 for depiction of the four methods for calculating 16S ratios and main text for description of treatment conditions.

### Diverse microbial taxa exhibit similar 16S RNA:DNA distributions

One consideration of using 16S ratios for activity estimations is that different taxa may have different thresholds of 16S ratios to qualify as ‘active’ (13). Choosing an arbitrary threshold 16S ratio for distinguishing ‘active’ from ‘inactive’ taxa therefore may be biased against taxa that have naturally low 16S ratios yet are truly active (and vice versa). To explore this possibility, we plotted the distributions of 16S ratios of the most abundant phyla, and pooled all less abundant phyla, in the corn and bean soil datasets. We found that in both soil types, all phyla exhibited a log-normal distribution of 16S ratios that were centered around a value of 1 (Fig. 4A and B). The finding that all phyla exhibited a similar distribution of 16s ratios suggests that altering the threshold 16S ratio for activity does not particularly impact specific taxa, at least at the phylum level. Therefore, our results support the use of a single threshold for 16S ratio activity analyses, even in diverse microbial communities.

**Fig 4.**
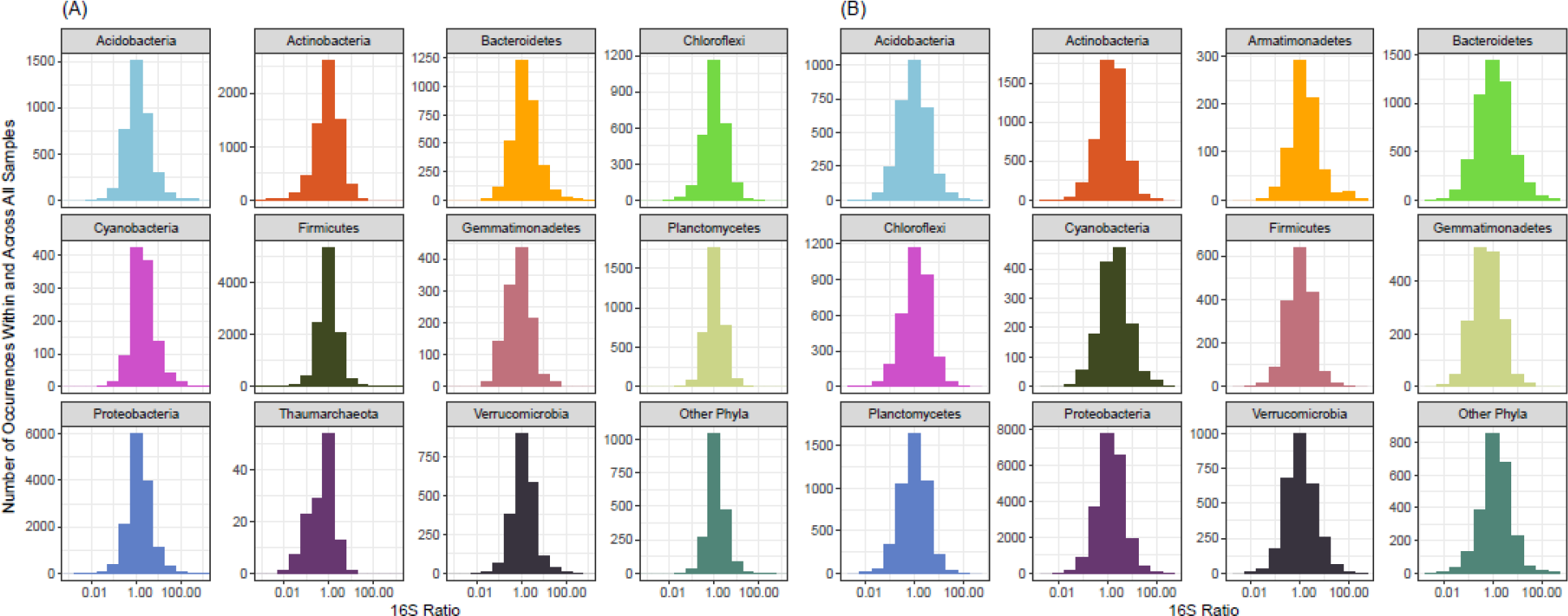
Histogram depicting the distribution of 16S rRNA:rDNA ratios for phyla in the rhizosphere soil of bean (A) and corn (B). The y-axis depicts the number of times a given 16S ratio occurs in the dataset (i.e. within and across all samples) for a given phylum. Note that the x-axis is on a log-scale. The most abundant phyla are shown, along with less abundant phyla pooled as ‘other phyla’.

An additional potential issue with using 16S ratios to estimate the proportion of active taxa community is the variability in copy numbers of the 16S rRNA operon across genomes of different taxa. 16S rRNA operon copy numbers can affect patterns of beta diversity in community structure (32) and can vary substantially between lineages (31). For example, lineages with many 16S operons (e.g. Firmicutes) may have lower 16S ratios because their abundance is overestimated by redundant OTUs in 16S rDNA data. To address this, we examined the relationship between the 16S ratio and the average number of ribosomal operons within phyla for all detected OTUs (Fig. 5). Although 16S ratios and average 16S rRNA operon count at the phylum level were correlated in both corn (r = −0.068, *p* < 0.0001) (Fig. 5A) and bean (r = −0.010, p = 0.0016) (Fig. 5B) rhizosphere soil, these correlations were extremely weak. In addition, across all operon counts, 16S ratios had similar ranges (Fig. 5). Recent work has advised against correcting for 16S rRNA operon counts in 16S rRNA gene surveys of microbial community structure, especially in taxa that are divergent from cultured representatives (32). Our results additionally suggest that accounting for 16S operon number likely has little effect on activity estimates in 16S ratio analyses.

**Fig 5.**
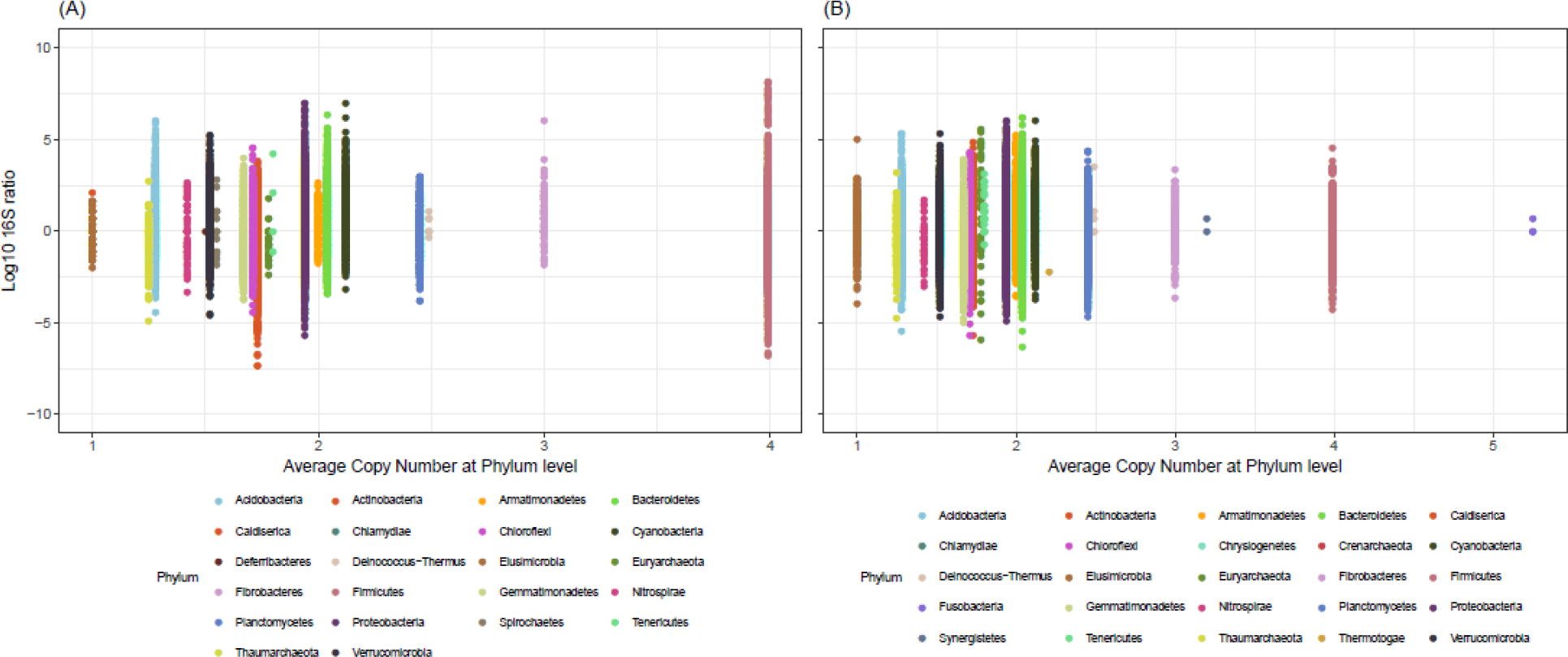
16S rRNA:rDNA ratio as a function of the average 16S operon copy number for the presented phyla as determined by the Ribosomal Copy Number Database (rrnDB). Data points represent every occurrence (i.e. within and across all samples) for a given phylum. Only phyla with representatives in the rrnDB are shown.

### CTC staining and 16S RNA:DNA capture complementary patterns of activity across treatments

Our final aim was to assess the level of agreement between two of the most common methods used for determination of microbial activity: the 16S ratio method and the CTC staining method. Across all treatments and between both methods, estimates of percent activity (i.e., between 10 and 60% of cells/taxa active) are similar to values reported in the literature for soil, as recently synthesized by (7). Though the two methods did not always agree in the level of percent activity detected, their overarching patterns across treatments were consistent with one exception.

In corn rhizosphere soil, the CTC staining method consistently resulted in higher estimates of proportional activity compared to the 16S ratio method. Using the CTC staining method, between 50-60% of the community was active for most treatments (all except the ‘post-dry’ treatment; Fig. 6A), while the 16S ratio method produced estimates in the range of 20-30% (Fig. 6C). Using both methods, percent activity sharply declined from the ‘pre-dry’ to the ‘post-dry’ treatment, and sharply increased in response to the ‘post-water’ treatment. This increase in response to watering was especially pronounced using the 16S ratio method, in which percent activity rebounded to a level which exceeded that of the pre-dried soil (Fig. 6C). After the ‘post-water’ treatment, percent activity increased in response to ABA, IAA, JA, and SA application using the CTC staining method, but did not change in response to the ‘control’ treatment (i.e., water alone) (Fig. 6A). In contrast, after the ‘post-water’ treatment, percent activity decreased in response to all four phytohormones, as well as in response to water-alone controls, using the 16S ratio method (Fig. 6C). In bean rhizosphere soil, the CTC staining method and the 16S ratio method produced more similar estimates of percent activity than for corn soil, and both methods produced identical patterns of percent activity across treatments (Fig. 6B and D) Using both methods, the drought and the nutrient addition treatments exhibited significantly lower percent activity compared to the control treatment, although the magnitude of this difference was lower using the 16S ratio method.

**Fig 6.**
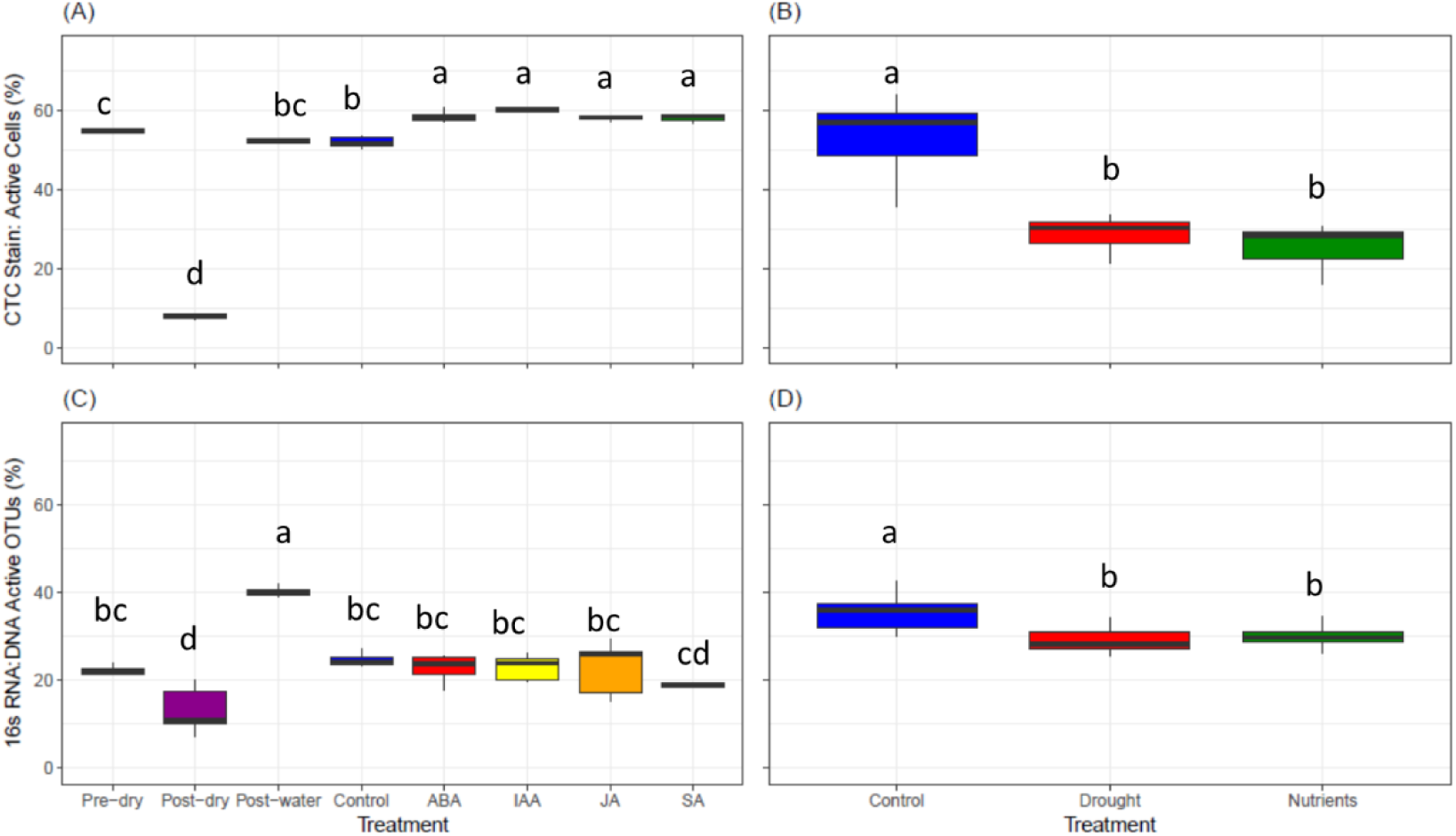
Proportion of active taxa/cells as determined by CTC/Syto24 staining (A, B) or 16s rRNA:rDNA ratios (C, D) in rhizosphere soil of corn (A, C) and bean (B, D). Taxa in (C, D) are defined as active with a 16S rRNA:rDNA ratio > 1. See main text for description of treatment conditions.

The extent of agreement between the 16S ratio method and the CTC staining method across the corn and bean soil experiments may be related to the duration of the experimental treatments in the two studies. In the corn soil experiment, the drying and re-wetting treatments lasts three and six days, respectively, while the phytohormone treatments lasted only 24 hours, and in the bean soil experiment, treatments were continuously applied for the approximately five weeks of the experiment. Given that differential transcription of RNA can occur within 30 seconds after an environmental stimulus (33) but cell doubling takes considerably longer (~10 minutes for only the very fastest-growing strains under optimal laboratory conditions (34), and many hours for others especially in natural environmental conditions (35 and references therein)), the 16S ratio method may be more sensitive in short time-frames than the CTC staining method. This may explain why the relatively short (24-hour) phytohormone treatments in corn soil resulted in relatively large decreases in percent activity using 16S ratios (average difference of 17.8% activity compared to ‘post-water’), but much smaller (albeit significant) increases in percent activity using CTC staining (average difference of 5.1% activity compared to ‘post-water’) (Fig. 6A and C). CTC staining, on the other hand, may perhaps give greater resolution in longer time-frames than 16S ratios. For example, the CTC staining method detected larger shifts in percent activity than the 16S ratio method in response to the three-day drying (46.1% decrease versus 9.1% decrease, respectively) and six-day re-wetting treatments (43.8% increase versus 27.1% increase, respectively) in corn rhizosphere soil (Fig. 6A and C), and in response to the five-week drought (23.1% decrease versus 6.3% decrease, respectively) and nutrient addition treatments (27.6% decrease versus 5.3% decrease, respectively) in bean soil (Fig. 6B and D). Despite differing in actual estimates of percent activity, both methods revealed very similar ecological patterns when considering treatments conducted over relatively long (≥ 3 days) periods, suggesting both methods provide robust metrics of whole community activity in experiments involving longer treatment exposures.

It is worth noting that the cell staining protocol used in our study (staining with both CTC and Syto24) allows for investigation of how both active and total cell counts varied across treatments and potentially influenced estimates of percent activity. In corn rhizosphere soil, both CTC counts (i.e., the number of active cells) and Syto24 counts (i.e., the total number of cells regardless of activity) sharply declined from the ‘pre-dry’ to the ‘post-dry’ treatment, and sharply increased in response to the ‘post-water’ treatment (Fig. S3A and C). After the ‘post-water’ treatment, both active and total cell counts decreased in response to ABA, IAA, JA, and SA application, as well as in response to the ‘control’ treatment (i.e., water alone). However, total cell counts were also significantly lower in the phytohormone treatments compared to the controls. This indicates that the increase in percent activity in response to phytohormones as revealed by cell activity staining (Fig. 6A) is due to a decrease in total cell counts coupled with an unchanging number of active cells. This may suggest potential toxicity of specific taxa in response to phytohormone application; either through direct phytohormone effects or indirect effects on soil pH. This is supported by the reduction in percent active OTUs in response to phytohormone application detected by the 16S method (Fig. 6C). In contrast, in bean rhizosphere soil, the decrease in percent activity reported by CTC staining due to drought and additional nutrients was due to a decrease in the number of active cells, as total cell counts were constant across treatments (Fig. S3B and D). This suggests an increase in microbial dormancy in response to water-limitation and ionic stress, potentially as a strategy to remain viable during sub-optimal environmental conditions.

### Considerations of the 16S ratio and CTC staining methods

An important consideration of the present work is that both the 16S ratio method and the CTC staining method have distinct biases that can influence percent activity estimates. A number of issues have been highlighted for analysis of bacterial activity with 16S ratios (13, 17, 36, 37), including the presence of dead cells and extracellular DNA, variations in the threshold ratio for bacterial activity, variations in sequencing depth, and molecular methodology (PCR biases). Although CTC staining avoids many assumptions of the 16S ratio method, it presents its own unique biases. For example, CTC staining in the present study excluded obligate anaerobes, potentially underestimating percent activity. In addition, CTC staining assumes that all (or at least a representative subsample) of the cells are extracted from the soil matrix, an assumption which may or may not be true, as reported for other studies performing cell counts from soil samples (21, 22). Finally, because CTC staining requires that the cells are extracted from the soil matrix prior to conducting the assay, it may introduce artefacts during the cell extraction process (i.e., cell death). Nevertheless, the finding of robust ecological patterns across long-term (≥ 3 days) treatments for the two methods, despite the different biases of each method, supports the conclusion that either method can provide a comparative assessment of community activity.

Another consideration of our analysis is the inability of the methods used here to account for both extracellular DNA and dead cells in the rhizosphere. Extracellular DNA (38, 39) and dead bacteria or necromass (40) are common across soils, and both can cause 16S ratios to underestimate percent activity (36). Similarly, although our CTC flow cytometry size gating likely excludes extracellular DNA by removing particles < 1 μm, intact dead cells would be included in the ‘total cell count’ calculated by Syto24 staining, thereby underestimating percent activity. Although our study was not designed to allow determination of extracellular DNA or dead cell abundance, we note that the rhizosphere is generally assumed to be an area of high metabolic activity. Thus, we might expect relic nucleic acids or dead cells to turn over relatively quickly, limiting their confounding effects in the present study. Our data support the rapid turnover of dead cells, given that the phytohormone treatments in the corn soil experiment, which lasted only 24 hours, led to significant and substantial decreases in total cell counts (Fig. S3C). It should also be noted that, in the corn soil experiment, soils were frozen prior to activity analyses, potentially increasing the number of dead cells and artificially reducing percent activity. We suggest this impact was minimal, given that percent activity estimates in the present study are relatively high compared to previous estimates in soil (7). Nevertheless, we suggest that combining our 16S ratio and CTC/Syto24 approach with a stain specific to extracellular DNA and dead/dying cells, such as propidium monoazide, would clarify the impact of these nucleic acid pools in assessments of microbial activity (41).

Finally, our study was conducted in rhizosphere soil, a very distinct environment relative to bulk soils and other natural environments due to the continual inputs of labile carbon by the plant host and high rates of metabolic activity. Expanding the environments where 16S ratios and activity staining are paired will inform on the general utility of the two methods for estimating microbial activity.

### Conclusions

Overall, our results provide insight into the estimation of microbial activity using two of the most commonly reported methods: 16S ratios and CTC staining. Although we found a high prevalence of phantom taxa in our sequencing datasets, patterns in percent activity across treatments were largely unaffected by the method used to account for these taxa. We also examined the potential for phylum-specific biases in 16S ratios, and found that 16S ratio distributions were highly similar across microbial phyla and were only weakly correlated with ribosomal operon number, suggesting that the use of a single 16S threshold does not introduce bias across broad phylogenetic groups. Lastly, we found that 16S ratios and CTC staining capture complementary patterns of activity across treatments. While the two methods revealed identical ecological patterns across longer-term (≥ 3 days) treatments, they produced an opposite pattern in response to shorter-term (24 hours) treatments; a finding likely related to the temporal resolution of the two methods. Our results suggest that 16S ratios and CTC staining provide robust and complementary estimates of bacterial and archaeal community activity in rhizosphere soils.

## Materials and Methods

We conducted two separate experiments (‘corn rhizosphere soil’ and ‘bean rhizosphere soil’), with varying stress treatments in each experiment. For clarity, we first present the methods specific to each experiment, and then present the methods shared between experiments.

### Corn rhizosphere soil: experimental design, sample collection, and preparation for sequencing

In the first experiment, topsoil was collected on August 21, 2017 from the AGR-Corn treatment of the Great Lakes Bioenergy Resource Center Scale-Up Experiment located near the Kellogg Biological Station, Hickory Corners, MI. Corn has been planted annually at that site since 2010. Replicate soil cores (to a depth of 10 cm) were collected using a 2.5 cm diameter steel corer, transported to the laboratory on ice, sieved and homogenized. Three replicate soil aliquots were weighed, dried for 72 hours at 70°C, then re-weighed to determine soil percent moisture (mean 8.6% ± 0.3% standard deviation), and the remaining soil was stored at 4°C until use.

Broadly, the experimental design consisted of several pre-treatments: ‘pre-dry’ (soil was sampled before any treatments were initiated), ‘post-dry’ (soil was dried and then sampled), and ‘post-water’ (soil was partially re-wetted and then sampled), before exposing soils to one of five treatments: application of the phytohormones abscisic acid (ABA), indole-3-acetic acid (IAA), jasmonic acid (JA) or salicylic acid (SA), or water control. On April 2, 2018, five replicates (30 g each) of the sieved and homogenized soil was retrieved from 4°C storage and frozen at −80°C for DNA/RNA extractions (i.e., the ‘pre-dry’ treatment). The remaining soil was dried for 72 hours at 45°C, at which point another five replicates (30 g each) were stored at −80°C (i.e., the ‘post-dry’ treatment). The remaining dried soil was split into 50 mL conical tubes (30 g of dry soil each), and each tube received water to achieve 4.3% percent soil moisture (half of the initial 8.6% percent soil moisture). This initial wetting step was included to isolate potential responses to phytohormones from the known response to moisture ((10) and references therein). Tubes were vigorously mixed and any clumps broken up with a sterile pipet. Tubes were incubated at room temperature for six days, then five replicates were frozen at −80°C (i.e., the ‘post-water’ treatment). The remaining tubes were then randomly assigned to one of five treatments: IAA, JA, ABA, SA, or water control. Five replicate tubes received 1.12 mL of the appropriate 0.22 μm filter-sterilized phytohormone dissolved in water at a concentration of 1 mM, while the control tubes received filter-sterilized water alone. Thus, these treatments restored all tubes to the initial 8.6% percent soil moisture. Tubes were vigorously mixed and clumps broken up with a sterile pipet. After 24 hours, the soil samples were frozen at −80°C.

DNA was extracted from ~0.23 g soil samples using the Qiagen PowerSoil kit following the manufacturers recommendations, while RNA was extracted from a protocol modified from (11, 42). Briefly, up to 0.5 g of soil was added to 200 μL of autoclaved PBL buffer (5 mM Na_2_-EDTA, 0.1% w/v sodium docecyl sulfate, and 5 mM Tris-HCl, pH ~3), vortexed for 1 minute, then 1 mL of phenol:chloroform:isoamyl alcohol (25:24:1 v/v/v, pH 8) was added. Samples were vortexed for 15 min, then centrifuged for 5 min at 20,000 x g. The upper (i.e., aqueous) layer was collected, added to 1 mL isopropanol, and vortexed. Samples were centrifuged for 15 min at 20,000 x g, and the supernatant was carefully removed. Tubes were air dried for 15 minutes, then resuspended in 50 μL of sterile water. Resuspended RNA extracts were cleaned using the OneStep PCR Inhibitor Removal kit (Zymo Research, Irvine, CA).

### Bean rhizosphere soil: experimental design, sample collection, and preparation for sequencing

In the second experiment, we planted 24 one-gallon pots with the common bean, *Phaseolus vulgaris* L. (var. Red Hawk), in local Michigan field soil in a controlled-environment growth chamber (BioChambers FXC-19). Plants received 16 h light and 8 h dark photoperiod, with a daytime temperature of 29°C and a nighttime temperature of 20°C. Eight replicate plants received ample water throughout the course of the experiment and served as controls. Eight additional replicates received ample water with the addition of nutrients (half-strength Hoagland solution; (43)) and an additional eight replicates were subjected to continuous drought, receiving 66% less water than control pots throughout the experiment. Plants were grown to the R8 stage (~5 weeks) before harvesting rhizosphere soils. Rhizosphere soil was collected in sterile Whirl-Pak bags by uprooting the plants and shaking loose soil from the root system. Any remaining soil adhering to the roots was considered rhizosphere soil. Two rhizosphere soil samples (5 g each) per plant were collected and immediately processed for active and total cell counts (see further detail below), while the remaining rhizosphere soil was frozen at −80°C for RNA/DNA extraction. For each plant, DNA was extracted from ~0.3 g of rhizosphere soil using the DNeasy Powersoil kit (Qiagen, Carlsbad, CA, USA, while RNA was extracted from ~2.3 g of rhizosphere soil using the RNeasy Powersoil kit, following manufacturer’s instructions.

### Corn and bean rhizosphere soil: microbial cell extraction and active and total cell counts

For both corn and bean rhizosphere soils, microbial cells were extracted following a protocol adapted from (44), and stained for determination of active and total cell counts. Briefly, rhizosphere soil (10 g per sample in the corn soil experiment, and two technical replicates of 5 g each in the bean soil experiment) was mixed with 100 ml of chilled sterile phosphate buffered saline containing 0.5% Tween-20 (PBST). Soil samples were homogenized in a Waring blender (Conair Corporation, East Windsor Township, NJ, USA) three times for one minute and kept on ice between each blending cycle. Soil slurries were centrifuged at 1,000 *x g* for 15 minutes and the supernatant was set aside. The pelleted soil was resuspended in 100 ml PBST and blended for an additional minute and re-centrifuged. The supernatants were pooled and centrifuged at 10,000 *x g* for 30 min. The supernatant was aspirated and the pellet was resuspended in sterile Milli-Q water (20 mL in the corn soil experiment, and 10 mL in the bean soil experiment). Cells were stained for percent activity determination using the BacLight RedoxSensor CTC Vitality kit (ThermoFisher Scientific, Waltham, MA, USA). Briefly, one milliliter of cells was stained with 0.38 μl of the DNA stain Syto24 and 5 mM of the activity stain 5-cyano-2,3-ditolyl tetrazolium chloride (CTC; active community) for 24 hours. Stained cells were fixed with 100 μl of 37% formaldehyde and cell counts were measured on a BD C6 Accuri Flow Cytometer (Franklin Lakes, NJ, USA), defining a cell as events which fluoresce above 10^3^ on a 490/515 nm filter for Syto24 and 450/630 nm filter for CTC. Following recommendations by the Michigan State University Flow Cytometry Core, we gated measurements by side scatter values <500 which removed particles <1 μm from our measurements.

We calculated the percentage of active cells by dividing CTC counts by Syto24 counts. For each sample in the corn soil experiment, we used a single 10 g sample of rhizosphere soil for microbial cell extraction that was then split into two technical replicates for staining: these two replicates per sample were averaged prior to subsequent analyses to avoid pseudoreplication. For each plant in the bean experiment, we used two replicate 5 g soil samples to give two technical replicate microbial cell extractions. Each of these was then split into three technical replicates for staining. These six replicates per plant were averaged prior to subsequent analyses to avoid pseudoreplication.

### Corn and bean rhizosphere soil: 16S rRNA gene amplicon sequencing

For both the corn and bean rhizosphere soil experiments, we first verified no DNA contamination in the RNA samples using PCR with 16S primers (45, 46) followed by gel electrophoresis. Next, 3 μl of RNA from each RNA sample was reverse transcribed using the SuperScript RT III kit (Invitrogen) following the protocol for random hexamers. Nucleic acid concentrations were measured with the Qubit broad-range DNA assay kit (ThermoFisher, Waltham, MA, USA). DNA and cDNA from the bean experiment were diluted to 5ng ul^−1^ (but were left undiluted in the corn soil experiment) prior to submitting for sequencing at the Michigan State Genomics Core. cDNA and DNA from both the corn and bean rhizosphere soil experiments were sequenced by the Michigan State University Genomics Core using the dual-indexed primer pair 515F and 806R (46). Samples were prepared for sequencing by the MSU Genomics Core including PCR amplification and library preparation using the Illumina TruSeq Nano DNA Library Preparation Kit. Paired-end, 250bp reads were generated on an Illumina MiSeq and the Genomics Core provided standard Illumina quality control, adaptor and barcode trimming, and sample demultiplexing.

### Corn and bean soil: bioinformatic and statistical analyses

The corn and bean rhizosphere soil sequencing datasets were analyzed separately. For each dataset, raw reads were merged, quality filtered, dereplicated, and clustered into 97% identity operational taxonomic units (OTUs) using the UPARSE pipeline (47). Taxonomic annotations for OTU representative sequences were assigned in the mothur (48) environment using the SILVA rRNA database release 132 (49). OTUs annotated as mitochondria or chloroplasts were removed, and OTU representative sequences were aligned using MUSCLE (version 3.8.31 (50)). Aligned sequences were used to build a phylogeny using FastTree (version 2.1.7 (51, 52)). The resulting tree was imported into R (version 3.3.2 (53)) using ‘ape’ (version 3.0.11 (54)) and was rooted at the midpoint using ‘phangorn’ (version 2.4.0 (55)). All subsequent analyses were performed in R (version 3.5.0), with ecological statistics performed using phyloseq (version 1.24.0 (56)). Data were visualized using a combination of the R packages ggplot2 (version 2.2.1; (57)), reshape2 (version 1.4.3; (58)), and gridExtra (version 2.3; (59)), and dplyr (version 0.7.5; (60)) was used for data summaries.

First, we examined the prevalence of ‘phantom taxa’ (i.e. OTUs with detectable RNA reads but no detectable DNA reads; (18)) in the corn and bean rhizosphere datasets. We calculated the average percent of OTUs that are phantom taxa in each treatment, as well as the average percent of OTUs with a single RNA read and no detectable DNA reads in each treatment. We conducted these analyses across a range of subsampling levels (using a step-size of 5000 reads per sample) in order to examine the influence of sequencing depth on the prevalence of phantom taxa, and used the loess smoothing function (61) to generate best fit lines and confidence intervals. Given the relatively low impact of subsampling level on the occurrence of phantom taxa, all subsequent analyses were conducted on samples rarefied to the minimum sampling depth in each dataset (22,589 reads per sample for corn soil, and 38,021 reads per sample for bean soil).

Given the prevalence and persistence (i.e., their high collective contributions regardless of sampling effort) of phantom taxa in our sequencing datasets, we next compared four different methods for calculating 16S ratios in the presence of phantom taxa. See Fig. 2 for a detailed illustration of the four methods, which we refer to as Methods 1, 2, 3, and 4 for simplicity. In Method 1, each phantom taxon in each sample is set to a 16S ratio of 100 in order to designate such taxa as ‘active’ regardless of the threshold 16S ratio activity level chosen, since most studies choose a threshold ratio less than 10 to designate taxa as ‘active’ (6, 8, 11). In addition, Method 1 sets each taxon in each sample with no detectable RNA or DNA reads to a value of zero, thereby eliminating undefined 16S ratios which arise due to a denominator of zero. In Method 2, every instance in which zero DNA reads are detected for a given OTU in a given sample is changed to a value of one in order to eliminate zeros in the denominator. In Method 3, previously used by (11), a value of one is added to every OTU in every sample in the DNA dataset. This method is meant to eliminate zeros in the denominator (as with Method 2), but also to treat every entry in the DNA dataset exactly the same. In Method 4, previously used by (14), a value of one is added to every OTU in every sample in both the RNA and the DNA datasets. As with Methods 2 and 3, this method is meant to eliminate zeros in the denominator, but also to treat every entry in the entire dataset (both RNA and DNA reads) exactly the same. We compared the resulting percent activity of the OTUs after using Methods 1-4 in both the corn and bean rhizosphere datasets, using threshold 16S ratios of 1, 2, and 5 for determination of ‘active’ versus ‘inactive’ OTUs. Given that Methods 1-4 captured similar patterns in percent activity across treatments, we conducted all subsequent analyses using the recently published Method 3 (11) to calculate 16S ratios.

Next, we examined phylum-level differences in the distribution of 16S ratios in both the corn and bean rhizosphere datasets. After calculating 16S ratios, we generated histograms of the number of times a given 16S ratio occurred in each dataset (i.e. including every OTU in every sample assigned to a given phylum). We also examined the relationship between the average number of 16S ribosomal operons per genome for each phylum, obtained from the Ribosomal RNA Database (version 5.4; (31, 62, 63)), and the observed 16S ratios in the present study.

Finally, we compared estimates and across-treatment patterns of microbial activity using the 16S ratio (threshold > 1) method to calculate the percentage of active taxa, versus using the cell staining method (CTC counts divided by Syto24 counts) to calculate the percentage of active cells. We also examined both active (CTC) and total (Syto24) counts across treatments in order to explore the influence these two values have on the percentage of active cells as calculated by the CTC/Syto24 ratio. Differences among treatments were assessed using ANOVA followed by a Tukey post-hoc test for multiple comparisons. All bioinformatic workflows and custom scripts are available on GitHub (https://github.com/ShadeLab/PAPER_Bowsher_16sRatio_CTCstain).

### Accession number(s)

Corn and bean rhizosphere sequencing data were submitted to the NCBI Sequence Read Archive under BioProject accession numbers PRJNA490178 and PRJNA454289, respectively.

## Acknowledgements

We would like to thank Jackson Sorensen, John Chodkowski, and Louis King for assistance and troubleshooting of flow cytometry methods and the Sheng Yang He lab for use of their flow cytometer.

## Funding

This work was supported in part by the Michigan State University Plant Resilience Institute, the National Science Foundation under grants DEB #1655425, DEB # #1749544, and MCB # #1817377, the USDA National Institute of Food and Agriculture, and Michigan State University through computational resources provided by the Institute for Cyber-Enabled Research. In addition, support for this research was provided by the U.S. Department of Energy, Office of Science, Office of Biological and Environmental Research (Awards DE-SC0018409 and DE-FC02-07ER64494), by the National Science Foundation Long-term Ecological Research Program (DEB 1637653) at the Kellogg Biological Station, and by Michigan State University AgBioResearch.

## Author Contribution

PJK and AS designed the experiments. PJK performed the experiments and all laboratory bench work. AB, PJK, and AS analyzed the data. AB wrote the paper, and all authors contributed to edits and revisions.

## Conflict of interest statement

The authors declare no conflict of interest.

---

**Fig S1.**
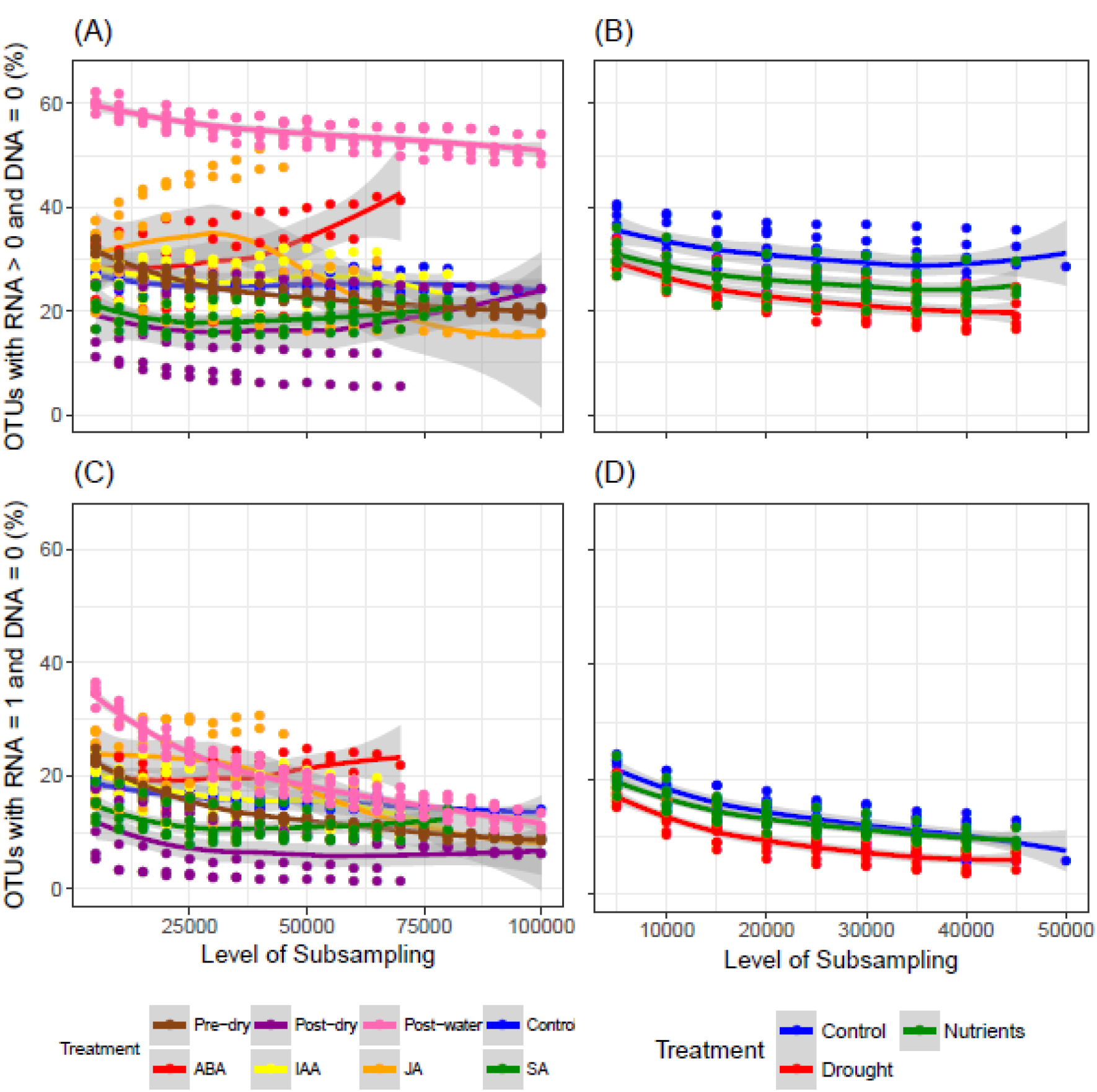
Prevalence of taxa with 16S RNA reads but zero 16S DNA reads (A, B) (i.e. ‘phantom taxa), or a single 16S RNA read and zero 16S DNA reads (C, D) in rhizosphere soil of corn (A,C) and bean (B,D) as a function of sequencing subsampling level. Points indicate individual samples, with best fit lines using the loess smoothing function. Gray shading around the smoothing lines are 95% confidence intervals. See Figure 1 for same plots but including only smoothing lines and confidence intervals.

**Fig S2.**
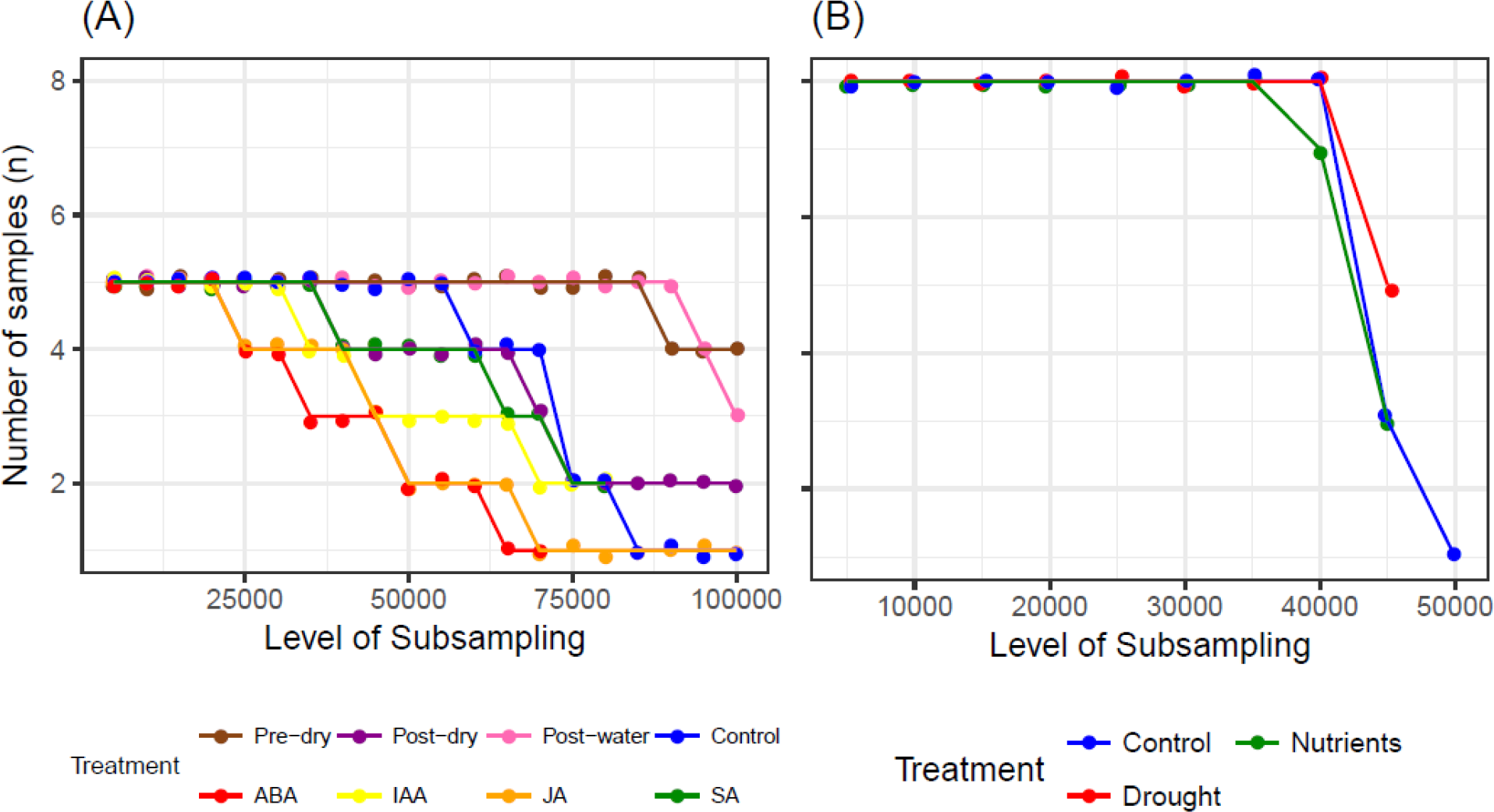
Number of samples in each treatment following subsampling to a given sequencing level in rhizosphere soil of corn (A, C) and bean (B, D). Note that the number of samples in each treatment decreases as subsampling level increases because samples are excluded when their total read count is less than the level of subsampling. See main text for description of treatment conditions.

**Fig S3.**
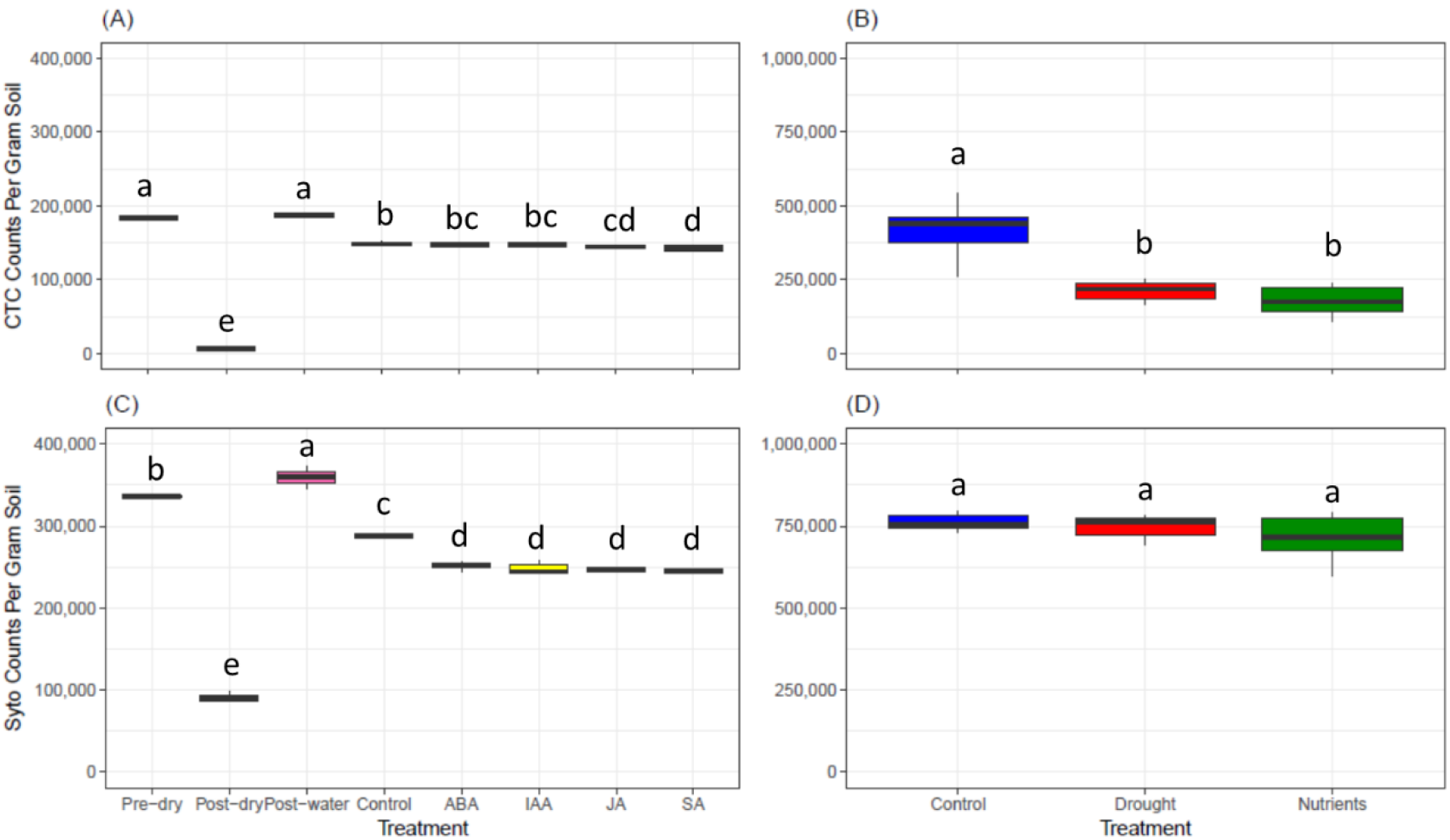
Flow cytometer counts (i.e., number of cells counted) per gram of soil extracted following staining with CTC (A, B) and Syto24 (C, D).

## References

1. Falkowski PG, Fenchel T, Delong EF. 2011. The microbial engines that drive Earth’s biogeochemical cycles. Science (80-) 320:1034–1039.

2. Arrigo KR. 2005. Marine microorganisms and global nutrient cycles. Nature 437:349–356.

3. Bardgett RD, Freeman C, Ostle NJ. 2008. Microbial contributions to climate change through carbon cycle feedbacks. ISME J 2:805–814.

4. Singh BK, Bardgett RD, Smith P, Reay DS. 2010. Microorganisms and climate change: terrestrial feedbacks and mitigation options. Nat Rev Microbiol 8:779–790.

5. van der Heijden MGA, Bardgett RD, van Straalen NM. 2008. The unseen majority: soil microbes as drivers of plant diversity and productivity in terrestrial ecosystems. Ecol Lett 11:296–310.

6. Blagodatskaya E, Kuzyakov Y. 2013. Active microorganisms in soil: Critical review of estimation criteria and approaches. Soil Biol Biochem 67:192–211.

7. Lennon JT, Jones SE. 2011. Microbial seed banks: The ecological and evolutionary implications of dormancy. Nat Rev Microbiol 9:119–130.

8. Jones SE, Lennon JT. 2010. Dormancy contributes to the maintenance of microbial diversity. Proc Natl Acad Sci 107:5881–5886.

9. Shoemaker WR, Lennon JT. 2018. Evolution with a seed bank: The population genetic consequences of microbial dormancy. Evol Appl 11:60–75.

10. Aanderud ZT, Jones SE, Fierer N, Lennon JT. 2015. Resuscitation of the rare biosphere contributes to pulses of ecosystem activity. Front Microbiol 6:1–11.

11. Kearns PJ, Angell JH, Howard EM, Deegan LA, Stanley RHR, Bowen JL. 2016. Nutrient enrichment induces dormancy and decreases diversity of active bacteria in salt marsh sediments. Nat Commun 7:1–9.

12. Wang G, Mayes MA, Gu L, Schadt CW. 2014. Representation of dormant and active microbial dynamics for ecosystem modeling. PLoS One 9.

13. Blazewicz SJ, Barnard RL, Daly RA, Firestone MK. 2013. Evaluating rRNA as an indicator of microbial activity in environmental communities: limitations and uses. ISME J 7:2061–2068.

14. Denef VJ, Fujimoto M, Berry MA, Schmidt ML. 2016. Seasonal succession leads to habitat-dependent differentiation in ribosomal RNA:DNA ratios among freshwater lake bacteria. Front Microbiol 7:1–13.

15. Franklin RB, Luria C, Ozaki LS, Bukaveckas PA. 2013. Community composition and activity state of estuarine bacterioplankton assessed using differential staining and metagenomic analysis of 16S rDNA and rRNA. Aquat Microb Ecol 69:247–261.

16. Aanderud ZT, Vert JC, Lennon JT, Magnusson TW, Breakwell DP, Harker AR. 2016. Bacterial dormancy is more prevalent in freshwater than hypersaline lakes. Front Microbiol 7:1–13.

17. Steven B, Hesse C, Soghigian J, Gallegos-Graves LV, Dunbar J. 2017. Simulated rRNA/DNA ratios show potential to misclassify active populations as dormant. Appl Environ Microbiol 83:1–11.

18. Klein AM, Bohannan BJM, Jaffe DA, Levin DA, Green JL. 2016. Molecular evidence for metabolically active bacteria in the atmosphere. Front Microbiol 7:1–11.

19. Shade A, Jones SE, Caporaso JG, Handelsman J, Knight R, Fierer N, Gilbert JA. 2014. Conditionally rare taxa disproportionately contribute to temporal changes in microbial diversity. MBio 5:1–9.

20. Hatzinger PB, Palmer P, Smith RL, Penarrieta CT, Yoshinari T. 2003. Applicability of tetrazolium salts for the measurement of respiratory activity and viability of groundwater bacteria. J Microbiol Methods 52:47–58.

21. Coyotzi S, Pratscher J, Murrell JC, Neufeld JD. 2016. Targeted metagenomics of active microbial populations with stable-isotope probing. Curr Opin Biotechnol 41:1–8.

22. Moran MA, Satinsky B, Gifford SM, Luo H, Rivers A, Chan LK, Meng J, Durham BP, Shen C, Varaljay VA, Smith CB, Yager PL, Hopkinson BM. 2013. Sizing up metatranscriptomics. ISME J 7:237–243.

23. Stellmach J. 1984. Fluorescent redox dyes I. Production of fluorescent formazan by unstimulated and phorbol ester-or digitonin-stimulated Ehrlich ascites tumor cells. Histochemistry 80:137–143.

24. Yamaguchi N, Nasu M. 1997. Flow cytometric analysis of bacterial respiratory and enzymatic activity in the natural aquatic environment. J Appl Microbiol 83:43–52.

25. Ullrich S, Karrasch B, Hoppe H-G, Jeskulke K, Mehrens M. 1996. Toxic effects on bacterial metabolism of the redox dye 5-cyano-2,3-ditolyl tetrazolium chloride. Appl Environ Microbiol 62:4587–4593.

26. Servais P, Agogué H, Courties C, Joux F, Lebaron P. 2001. Are the actively respiring cells (CTC+) those responsible for bacterial production in aquatic environments? FEMS Microbiol Ecol 35:171–179.

27. Wilhelm L, Besemer K, Fasching C, Urich T, Singer GA, Quince C, Battin TJ. 2014. Rare but active taxa contribute to community dynamics of benthic biofilms in glacier-fed streams. Environ Microbiol 16:2514–2524.

28. Houlden A, Timms-Wilson TM, Day MJ, Bailey MJ. 2008. Influence of plant developmental stage on microbial community structure and activity in the rhizosphere of three field crops. FEMS Microbiol Ecol 65:193–201.

29. Berg G, Smalla K. 2009. Plant species and soil type cooperatively shape the structure and function of microbial communities in the rhizosphere. FEMS Microbiol Ecol 68:1–13.

30. Vives-Peris V, Gómez-Cadenas A, Pérez-Clemente RM. 2017. Citrus plants exude proline and phytohormones under abiotic stress conditions. Plant Cell Rep 36:1971–1984.

31. Lee ZMP, Bussema C, Schmidt TM. 2009. rrnDB: Documenting the number of rRNA and tRNA genes in bacteria and archaea. Nucleic Acids Res 37:489–493.

32. Louca S, Doebeli M, Parfrey LW. 2018. Correcting for 16S rRNA gene copy numbers in microbiome surveys remains an unsolved problem. Microbiome 6:1–12.

33. von Wulffen J, Ulmer A, Jäger G, Sawodny O, Feuer R. 2017. Rapid sampling of Escherichia coli after changing oxygen conditions reveals transcriptional dynamics. Genes (Basel) 8.

34. Eagon RG. 1962. Pseudomonas natriegens, a marine bacterium with a generation time of less than 10 minutes. J Bacteriol 83:736–737.

35. Gibson B, Wilson DJ, Feil E, Eyre-Walker A. 2018. The distribution of bacterial doubling times in the wild. Proc R Soc B Biol Sci 285.

36. Dlott G, Maul JE, Buyer J, Yarwood S. 2015. Microbial rRNA:RDNA gene ratios may be unexpectedly low due to extracellular DNA preservation in soils. J Microbiol Methods 115:112–120.

37. Papp K, Hungate BA, Schwartz E. 2018. Comparison of microbial ribosomal RNA synthesis and growth through quantitative stable isotope probing with H218O. Appl Environ Microbiol 84:AEM.02441–17.

38. Carini P, Marsden PJ, Leff JW, Morgan EE, Strickland MS, Fierer N. 2016. Relic DNA is abundant in soil and obscures estimates of soil microbial diversity. Nat Microbiol 2:1–6.

39. Levy-Booth DJ, Campbell RG, Gulden RH, Hart MM, Powell JR, Klironomos JN, Pauls KP, Swanton CJ, Trevors JT, Dunfield KE. 2007. Cycling of extracellular DNA in the soil environment. Soil Biol Biochem 39:2977–2991.

40. Kallenbach CM, Grandy AS, Frey SD, Diefendorf AF. 2015. Microbial physiology and necromass regulate agricultural soil carbon accumulation. Soil Biol Biochem 91:279–290.

41. Emerson JB, Adams RI, Román CMB, Brooks B, Coil DA, Dahlhausen K, Ganz HH, Hartmann EM, Hsu T, Justice NB, Paulino-Lima IG, Luongo JC, Lymperopoulou DS, Gomez-Silvan C, Rothschild-Mancinelli B, Balk M, Huttenhower C, Nocker A, Vaishampayan P, Rothschild LJ. 2017. Schrödinger’s microbes: Tools for distinguishing the living from the dead in microbial ecosystems. Microbiome 5:86.

42. Mettel C, Kim Y, Shrestha PM, Liesack W. 2010. Extraction of mRNA from soil. Appl Environ Microbiol 76:5995–6000.

43. Hoagland DR, Arnon DI. 1950. The water-culture method for growing plants without soil. California Agricultural Experiment Station, Circular 347.

44. Portillo MC, Leff JW, Lauber CL, Fierer N. 2013. Cell size distributions of soil bacterial and archaeal taxa. Appl Environ Microbiol 79:7610–7617.

45. Walters W, Hyde ER, Berg-Lyons D, Ackermann G, Humphrey G, Parada A, Gilbert JA, Jansson JK, Caporaso JG, Fuhrman JA, Apprill A, Knight R. 2015. Improved bacterial 16S rRNA gene (V4 and V4-5) and fungal internal transcribed spacer marker gene primers for microbial community surveys. mSystems 1:1–10.

46. Kozich JJ, Westcott SL, Baxter NT, Highlander SK, Schloss PD. 2013. Development of a dual-index sequencing strategy and curation pipeline for analyzing amplicon sequence data on the MiSeq Illumina sequencing platform. Appl Environ Microbiol 79:5112–5120.

47. Edgar RC. 2013. UPARSE: highly accurate OTU sequences from microbial amplicon reads. Nat Methods 10:996–1000.

48. Schloss PD, Westcott SL, Ryabin T, Hall JR, Hartmann M, Hollister EB, Lesniewski RA, Oakley BB, Parks DH, Robinson CJ, Sahl JW, Stres B, Thallinger GG, Van Horn DJ, Weber CF. 2009. Introducing mothur: open-source, platform-independent, community-supported software for describing and comparing microbial communities. Appl Environ Microbiol 75:7537–7541.

49. Quast C, Pruesse E, Yilmaz P, Gerken J, Schweer T, Yarza P, Peplies J, Glockner FO. 2013. The SILVA ribosomal RNA gene database project: improved data processing and web-based tools. Nucleic Acids Res 41:590–596.

50. Edgar RC. 2004. MUSCLE: multiple sequence alignment with high accuracy and high throughput. Nucleic Acids Res 32:1792–1797.

51. Price MN, Dehal PS, Arkin AP. 2009. FastTree: computing large minimum evolution trees with profiles instead of a distance matrix. Mol Biol Evol 26:1641–1650.

52. Price MN, Dehal PS, Arkin AP. 2010. FastTree 2 – Approximately maximum-likelihood trees for large alignments. PLoS One 5.

53. Team RC. 2018. R: a language and environment for statistical computing. R Foundation for Statistical Computing, Vienna, Austria.

54. Paradis E, Claude J, Strimmer K. 2004. APE: analyses of phylogenetics and evolution in R language. Bioinformatics 20:289–290.

55. Schliep KP. 2011. phangorn: phylogenetic analysis in R. Bioinformatics 27:592–593.

56. Mcmurdie PJ, Holmes S. 2013. phyloseq: An R package for reproducible interactive analysis and graphics of microbiome census data. PLoS One 8.

57. Wickham H. 2009. ggplot2: Elegant graphics for data analysis. Springer-Verlag, New York.

58. Wickham H. 2007. Reshaping data with the reshape package. J Stat Softwawre 21.

59. Auguie B. 2017. gridExtra: miscellaneous functions for “grid” graphics. R package version 2.3.

60. Wickham H, Francois R, Henry L, Muller K. 2018. dplyr: a grammar of data manipulation. R package version 0.7.5.

61. Cleveland WS. 1979. Robust locally weighted regression and smoothing scatterplots. J Am Stat Assoc 74:829–836.

62. Stoddard SF, Smith BJ, Hein R, Roller BRK, Schmidt T. 2015. rrnDB: improved tools for interpreting rRNA gene abundance in bacteria and archaea and a new foundation for future development. Nucleic Acids Res 43:593–598.

63. Klappenbach JA, Saxman PR, Cole JR, Schmidt TM. 2001. rrndb: the ribosomal RNA operon copy number database. Nucleic Acids Res 29:181–184.

